# Predicting efficacy of immunotherapy in mice with triple negative breast cancer using a cholesterol PET radiotracer

**DOI:** 10.1101/2023.10.02.560577

**Authors:** Nicholas G. Ciavattone, Jenny Guan, Alex Farfel, Timothy Desmond, Benjamin L. Viglianti, Peter JH Scott, Allen F. Brooks, Gary D. Luker

## Abstract

Predicting the response to cancer immunotherapy remains an unmet challenge in triple-negative breast cancer (TNBC) and other malignancies. T cells, the major target of current checkpoint inhibitor immunotherapies, accumulate cholesterol during activation to support proliferation and signaling. The requirement of cholesterol for anti-tumor functions of T cells led us to hypothesize that quantifying cellular accumulation of this molecule could distinguish successful from ineffective checkpoint immunotherapy. To analyze accumulation of cholesterol by T cells in the immune microenvironment of breast cancer, we leveraged a novel positron emission tomography (PET) radiotracer, FNP-59. FNP-59 is an analog of cholesterol that our group has validated as an imaging biomarker for cholesterol uptake in pre-clinical models and initial human studies. In immunocompetent mouse models of TNBC, we found that elevated uptake of exogenous labeled cholesterol analogs functions as a marker for T cell activation. When comparing immune checkpoint inhibitor (ICI)-responsive EO771 tumors to non-responsive AT-3 tumors, we found significantly higher uptake of a fluorescent cholesterol analog in T cells of the ICI-responsive tumors both in vitro and in vivo. Using the FNP-59 radiotracer, we discovered that accumulation of cholesterol by T cells increased further in ICI-responding tumors that received ant-PD-1 checkpoint immunotherapy. We verified these data by mining single cell sequencing data from patients with TNBC. Patients with tumors containing cycling T cells showed gene expression signatures of cholesterol uptake and trafficking. These results suggest that uptake of exogenous cholesterol analogs by tumor-infiltrating T cells predict T cell activation and success of ICI therapy.

## Introduction

Immune checkpoint inhibitor antibodies (ICIs) blocking the PD-1 or CTLA-4 pathways have revolutionized cancer therapy. By preventing signals that turn “off” killing functions of T cells, ICIs can unleash host immunity against cancer cells, producing remarkable survival advantages and even cures for a minority of patients. Unfortunately, the majority of patients do not benefit from ICIs and some patients experience the severe side effects of these drugs. Extending ICIs to breast cancer and other, less immunogenic tumors remains a substantial barrier in cancer immunotherapy. Recently, the anti-PD-1 ICI pembrolizumab received approval for treatment of high-risk, early-stage triple negative breast cancer (TNBC) in July, 2021 (1). TNBC tumors, the least common and most aggressive subtype, typically show more inflammation than other subtypes of breast cancer. An inflamed, immune “hot” environment correlates with better outcomes in TNBC (2). Unfortunately, tumors in most patients with TNBC are not conducive to immunotherapy based on current metrics for selection and/or do not respond to ICIs (3). For example, in the KEYNOTE-355 trial, pembrolizumab significantly extended survival in patients categorized as favorable for immunotherapy based on high expression of PD-L1 in tumor cells, lymphocytes, and macrophages in biopsy samples (4). Despite improved overall survival, anti-PD-1 treatment allowed only 11% more patients to survive through the 4.5-year study endpoint relative to those treated only with chemotherapy. These data highlight shortcomings of existing criteria to predict patients likely to respond to immunotherapy in breast cancer. Creating new, optimally non-invasive methods to improve selection of patients for immunotherapy in TNBC and identify early effects of therapy on T cells will help optimize use of existing drugs and facilitate translation of new immunotherapies. Since only a minority of patients across all malignancies respond to current ICIs, better methods to select patients and monitor therapy likely will improve outcomes across multiple cancers.

Activated T cells are one of the hallmarks of an inflamed, effective anti-tumor immune response. Previous studies suggest a link between activation of T cells and cholesterol. T cells increase cholesterol through de novo synthesis from acetyl CoA or uptake from lipoproteins through receptor-mediated endocytosis of low-density lipoprotein (LDL) particles from the bloodstream. Cholesterol is an essential component of cellular membranes, where this lipid modulates membrane fluidity, permeability, and receptor mediated signaling(5). Upon activation, T-cell receptor (TCR) nanoclusters form in cholesterol-rich membrane domains to elicit strong TCR signaling. Proliferating T cells and other cell types require increased amounts of cholesterol to replicate cell membranes, utilizing cholesterol stored as cholesterol esters in intracellular lipid droplets. Despite requirements for cholesterol in T cell activation and proliferation, increased cholesterol uptake by tumor infiltrating T cells has been linked to an “exhausted” T cell phenotype(6). However, exhausted T cells indicate a history of active T cell populations, and both populations of T cells correlate with success of ICIs in TNBC (7, 8). Therefore, cholesterol uptake by tumor infiltrating lymphocytes could indicate a “hot” tumor environment and predict the efficacy of immunotherapy.

Our research group previously developed the fluorinated radiotracer, 18F-FNP-59 or FNP-59, as an analog for cholesterol that can be leveraged to detect cholesterol uptake by cells and tissues using positron emission tomography (PET)(9). FNP-59 overcomes limitations of previous generations of this radiotracer that have been used clinically to detect adrenal cortical tumors, while maintaining sensitivity to detect uptake in tissues and cells, including cells of the immune system(9). As a cholesterol analog, FNP-59 distributes into the lipid pool and enters lipoproteins with subsequent trafficking into cells through receptors like the low-density lipoprotein receptor (LDLR) and fatty acid transporter, CD36. The fluorine-18 radiolabel is used commonly in radiotracers for PET imaging studies, such as the glucose analog 2-[(18)F] fluoro-2-deoxy-D-glucose to detect glycolytic metabolism in cancer and other diseases(10, 11). We hypothesized that activated and primed T cells in the tumor microenvironment have increased uptake of cholesterol detectable with FNP-59 as a marker for success of ICI therapy.

This study reveals that increased uptake of cholesterol marks activation of T cells in mouse TNBC tumors responsive to ICI therapy, and we can distinguish T cells from ICI-responsive and non-responsive tumors with FNP-59. Ex vivo restimulation of T cells recovered from ICI responsive tumors in mice exhibited greater cholesterol uptake compared to non-responsive tumors. *In vivo,* we discovered that T cells from responsive tumors accumulated significantly more cholesterol than non-responsive tumors, the latter of which exhibited almost undetectable uptake. Lastly, data mining studies of human TNBC showed upregulation of genes-related to the uptake and trafficking of cholesterol in populations of activated T cells. These data demonstrate that uptake of cholesterol as measured with FNP-59 demarks proliferating T cell populations and correlates with success of immunotherapy in pre-clinical mouse models of TNBC. Moreover, our work points to FNP-59 as a potential imaging probe to improve monitoring of cancer immunotherapy.

## Results

### T cells from ICI responsive tumors show greater cholesterol uptake

To study cholesterol uptake and the immunotherapy response in tumor infiltrating T cells, we utilized cell lines that model TNBC, but exhibit different responses to immunotherapy. EO771 cells, one of the only mammary tumor cell lines derived from spontaneous tumorigenesis in C57BL/6J mice, responds to immunotherapy and has been used to test effects of therapeutic agents to improve immunogenicity(12). In contrast, AT-3 cells are derived from transgenic MMTV-PyMT mice and respond poorly to immunotherapies, including anti-PD-1, despite the presence of intertumoral T cells(13, 14). To directly compare effects of anti-PD-1 antibody therapy on tumor growth, we orthotopically implanted either EO771 or AT-3 breast cancer cells with syngeneic immortalized mouse mammary fibroblasts. Three days after implanting cells, we randomly assigned mice to treatment with anti-PD-1 antibody therapy or vehicle. Mice received a total of four doses administered every 3 days. We quantified tumor size by caliper measurements and monitored survival until mice reached humane endpoints for tumor burden. Mice with EO771 tumors showed remarkable reductions in tumor growth (Figure 1A & 1B) and significantly increased overall survival in response to anti-PD-1 therapy (Figure 1C). By comparison, anti-PD-1 treatment elicited no change in tumor growth (Figure 1D & 1E) or survival over control in mice with AT-3 tumors (Figure 1F). These data validate responsive and treatment refractory mouse models of immunotherapy in TNBC, providing contrasting systems to investigate cholesterol as a marker of T cell activation.

**Figure 1.**
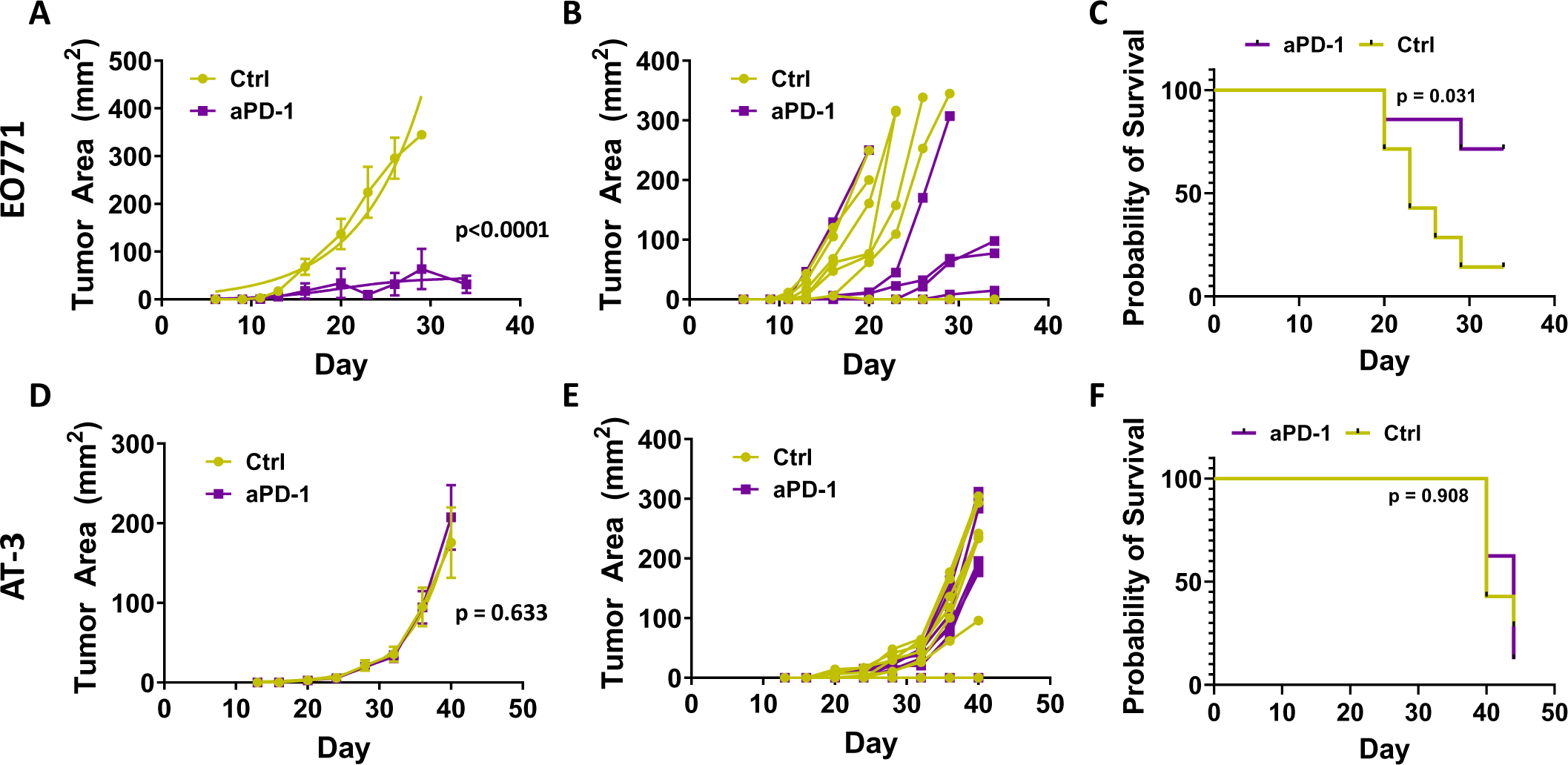
EO771 tumors respond to anti-PD-1 immunotherapy while AT-3 tumors do not. Three days after orthotopically injecting EO771 or AT-3 breast cancer cells plus mouse mammary fibroblasts into syngeneic C57BL/6j mice, we randomly assigned animals to treatment with anti-PD-1 antibody or isotype/vehicle ctrl every 3 days for 4 doses total. Graphs show mean values ± SEM (symbols) and calculated logistic regression (smooth line) for EO771 (A) or AT-3 (D) tumors (n=5 per group) treated with anti-PD-1 antibody or ctrl. Panels B (EO771) and E (AT-3) show tumor growth for individual mice over time. We analyzed differences in tumor growth data by logistic regression. Survival curves demonstrate that anti-PD-1 treatment significantly prolonged survival for mice with EO771 tumors (C) but not AT-3 (F) as analyzed by a Mantel-Cox test.

We measured uptake of cholesterol in T cells isolated from EO771 or AT-3 breast tumors and restimulated ex vivo in the presence of fluorescent 3-NBD-cholesterol. Due to the location and orientation of the NBD tag, this fluorescent cholesterol models cholesterol orientation in membranes better than previous fluorescently tagged cholesterol analogs (15). Solid tumors rarely contain naïve T cells thus ex-vivo tumor infiltrating T cells were stimulated with low level CD3ε for a brief period for re-stimulation(16). After re-stimulation, we assessed uptake of 3-NBD-Cholesterol in combination with activation marker CD69 on T cells using spectral cytometry (Figure 2A & 2D). Cholesterol uptake correlated significantly with expression of the activation marker, CD69 (Figure 2B & 2C). Additionally, cholesterol uptake increased to a markedly greater extent in CD8+ and CD4+ T cells from EO771 tumors when compared to T cells from AT-3 (non-responsive) tumors (Figure 2E&F).

**Figure 2.**
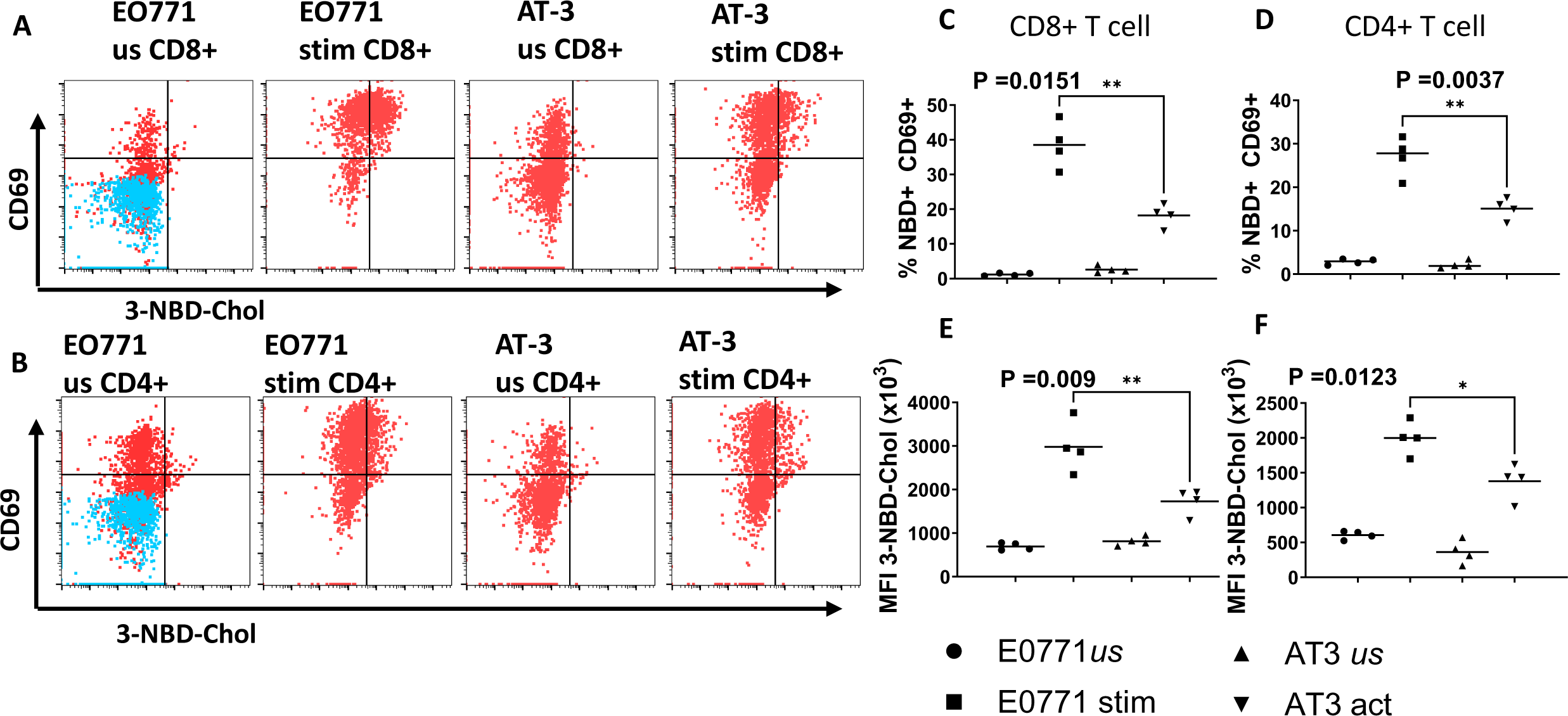
Greater ex vivo uptake of cholesterol in T cells from ICI responsive tumors. We isolated T cells from ICI responsive EO771 or unresponsive AT-3 tumors and left cells unstimulated (us) or re-stimulated on anti-CD3e coated (stim) dishes for 18 hours in the presence of 3-NBD labeled cholesterol. We determined uptake of labeled cholesterol in activated, CD69+ CD8 (A) or CD4 (B) T cells by flow cytometry. CD8 (C) and CD4 (D) T cells from EO771 tumors show significantly greater percentages of CD69+ cells with uptake of 3-NBD cholesterol than from ICI non-responsive AT-3 tumors. Stimulated CD8 (E) and CD4 (F) T cells from EO771 tumor also exhibited significantly higher mean fluorescence intensity for cholesterol uptake. Data are combined from 2 experiments with statistical comparisons by Student’s T test.

To assess uptake of cholesterol in T cells *in vivo*, we injected tumor bearing mice with a fluorescent BODIPY labeled cholesterol. BODIPY labeled cholesterol effectively labels intracellular cholesterol; remains stable in vivo; and exhibits greater fluorescence than the 3-NBD label(17). Twenty-four hours after injecting BODIPY-cholesterol, we collected intra-tumoral T cells for flow cytometry. T cells from responsive EO771 tumors showed a higher uptake of labeled cholesterol when compared to AT-3 tumors as seen in histograms (Figure 3A & 3B) and normalized BODIPY MFI in both CD8+ and CD4+ T cells (Figure 3C & 3D). Prior studies reported increased expression of PD-1 on T cells that take up more cholesterol. Indeed, PD-1 expression on T cells increased significantly on CD8+ T cells from EO771 tumors with higher cholesterol uptake (Figure 3E). Modulation of PD-1 did not occur on CD4+ T cells (Figure 3F). Together these experiments demonstrate increased uptake of cholesterol in T cells from tumors that respond to immunotherapy.

**Figure 3.**
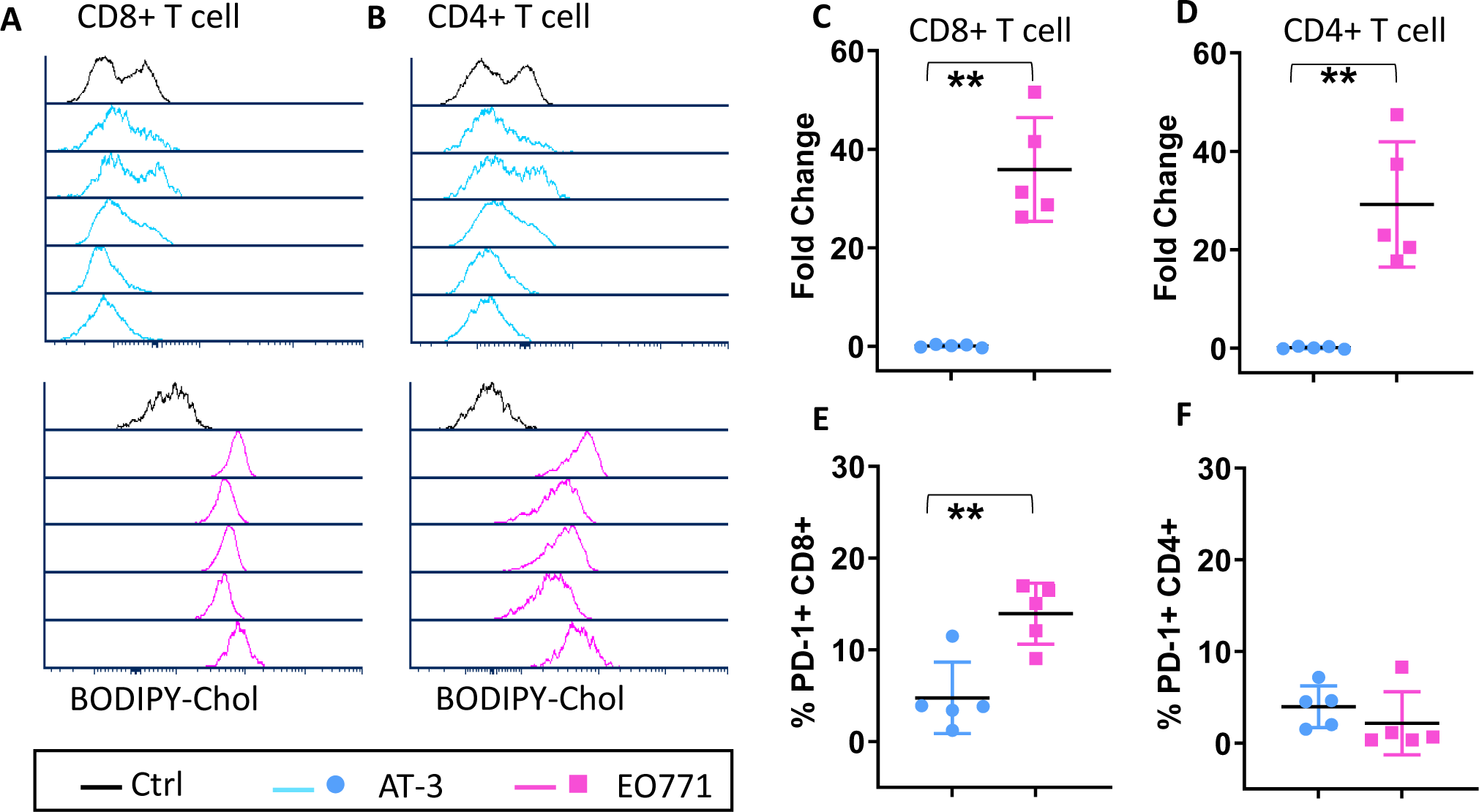
T cells in ICI responsive EO771 tumors show greater uptake of cholesterol in vivo. We injected C57BL/6J mice intraperitoneally with cholesterol labeled with red-shifted BODIPY and euthanized animals 24 hours later to collect and dissociate tumors for flow cytometry (n=5 each for EO771 and AT-3). Plots for (A) CD8+ and (B) CD4+ T cells show accumulation of BODIPY-cholesterol in cells from individual tumors from EO771 and AT-3 tumors relative to veh only/isotype antibody control. (C) CD8 and (D) CD4 T cells in EO771 tumors showed significantly higher fold change accumulation of fluorescent cholesterol relative to FMO control. (E) CD8 but not (F) CD4 T cells in EO771 tumors also expressed higher levels of PD-1. We compared differences between means using nonparametric Mann-Whitney tests (n = 5 mice per group) while differences between T cell population percentages were tested using Student’s T test.

### Activated T cells increase cholesterol uptake

Cholesterol is required for cell division processes, and its presence in the plasma membrane of T cells stabilizes TCR nanoclusters to enhance overall activation(18). Cholesterol uptake in T cells remains a controversial topic. One study found that hypercholesteremia in T cells reflected proliferation of T cells.(19) In contrast, elevated cholesterol in T cells also reportedly reflects exhaustion (6). To further investigate to what extent levels of cholesterol change with activation, we treated naïve mouse T cells with increasing amounts of anti-CD3ε and proportional increases in anti-CD28 antibodies (3 times the anti-CD3ε concentration). Using an in vitro assay to quantify total cholesterol, free cholesterol, and cholesterol esters, we established that activation increased total cholesterol concentration in T cells with differences between 0µg/mL and 1 µg /mL or 10 µg /mL of anti-CD3e (Figure 4A). The concentration of cholesterol esters did not change with increasing activation in T cells after 24 hours (Figure 4A). While this assay demonstrates that total cholesterol increased in T cells after TCR activation, these data do not distinguish between synthesis and uptake of cholesterol.

**Figure 4.**
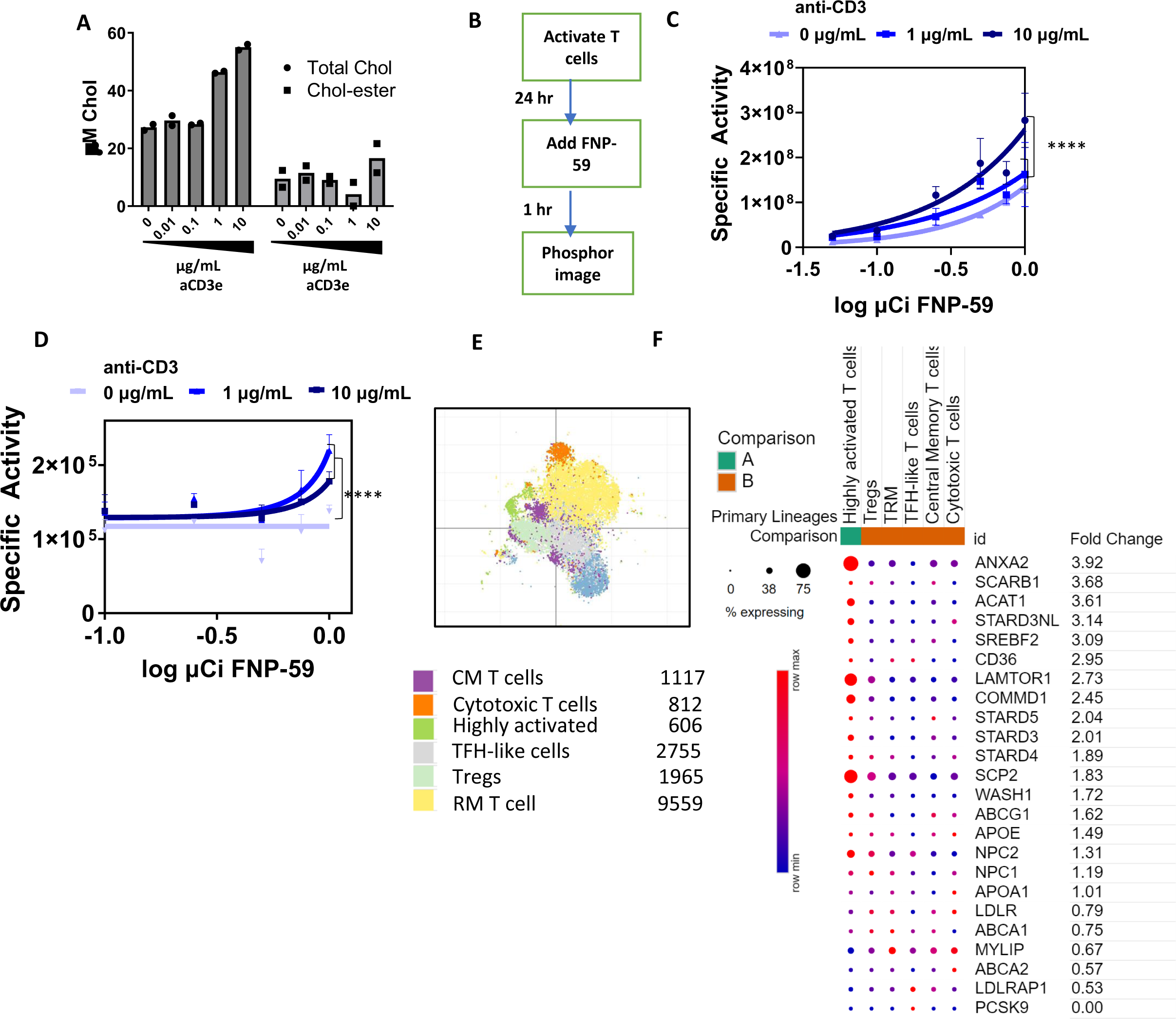
Cholesterol analog PET radiotracer, FNP-59, reveals increased uptake of cholesterol in activated T cells. (A) To check total cholesterol in T cells, we activated naïve mouse T cells with increasing concentrations of anti-CD3e for 24 hours before quantifying total and esterified cholesterol with a bioluminescence assay. Graphs show mean for total cholesterol and cholesterol esters (n=2 per condition; repeated twice). (B) We activated T cells for 24 hours prior to adding different concentrations of the cholesterol radiotracer analog, FNP-59, for one hour before washing and quantifying cell-associated radioactivity by autoradiography with a phosphor imaging screen. (C) Quantified data for uptake of FNP-59 at different amounts of administered radiotracer. We performed all experiments in triplicate, using comparison of fits with the extra sum-of-squares F test used to compare and test each data set and EC50 against the null hypothesis. Additionally, (D) T cells isolated from healthy human PBMC were activated with human anti-CD3e and treated and analyzed as described for mouse T cells. (E) Using annotated T cell clusters from the Immunological Genome Project’s (Immgen) Immune Cell Atlas scRNA-seq data sets on healthy human lamina propria [19], we analyzed genes in T cells involved in cholesterol uptake, retention, and trafficking. (F) Panel displays fold change in selected genes related to cholesterol metabolism between the ‘highly activated T cell’ cluster as compared with all other T cells.

To selectively measure uptake of cholesterol during T cell activation, we leveraged a novel PET radiotracer analog of cholesterol, FNP-59. Our group has validated FNP-59 as an imaging probe for cholesterol trafficking and uptake in humans(9). Since we added FNP-59 to the culture medium, any accumulated radiotracer in T cells represents uptake, rather than de novo synthesis, during activation. We incubated activated T cells with FNP-59 according to the diagram in Figure 4B. After washing cells to remove extracellular radiotracer, we used autoradiography with a phosphor imaging screen to measure uptake by T cells. After one hour of incubation, uptake of FNP-59 increased progressively with higher concentrations of activating antibodies. Each T cell treatment displayed significant difference between levels of anti-CD3 stimulation with unique EC50 values when tested against all data points in logistical nonlinear regression (Figure 4C).

To extend these findings to humans, we treated activated human T cells from peripheral blood with FNP-59 as outlined in figure 4B. We found human T cells also take up FNP-59 when activated, but higher activation of anti-CD3 was stymied when compared to mice (Figure 4D). To study further the impact of cholesterol trafficking in the human immune system we analyzed data from the Immune Cell Atlas, originating from Martin, et. al. 2019 and hosted by singlecell.broadinstitute.org(20). These data comprise an annotated set of immune cells from an uninflamed lamina propria, an excellent tissue site for studying varied immune responses without the presence of disease. From this dataset, we utilized the T cell clusters previously annotated by gene expression (Figure 4E). Genes responsible for uptake and trafficking of cholesterol displayed increased relative expression in the ‘Cycling’ cluster compared to other clusters (Figure 4F). These included genes specifically annotated for cholesterol uptake and trafficking, including scavenger receptors like SCP2 and SCARB1, endosome associated proteins such as STARD3/NL, NPC1/2, and transcription factor SREBF2(21–25). Additional upregulated genes, COMMD1, WASH1, ANXA2, and LAMTOR1, function in cholesterol uptake and trafficking as well as other cellular processes (26–29). Increased expression of genes related to cholesterol uptake and trafficking in activated human T cells supports translatability of these findings.

### Uptake of FNP-59 in T cells predicts and monitors response to immunotherapy in mouse TNBC

To test uptake of FNP-59 in T cells in vivo and use of this radiotracer to predict and monitor response to immunotherapy, we established mice with EO771 or AT-3 breast tumors. After tumors reached ∼ 1 cm diameter, we randomly assigned mice to treatment with anti-PD1 or vehicle. We hypothesized that anti-PD-1 treatment would further alter cholesterol uptake with reinvigoration of endogenous T cells. Four days later, we intravenously injected mice with 100µCi of FNP-59. We quantified uptake of FNP-59 in total dissociated splenocytes, and T cells recovered from dissociated tumors. Splenocytes from mice treated with vehicle only accumulated comparable amounts of FNP-59 for both EO771 or AT-3 groups (Figure 5A). Relative to vehicle only, anti-PD1 treatment modestly increased radiotracer uptake in splenocytes from both groups, although amounts did not differ significantly. Isolated T cells from the vehicle treated EO771 tumors accumulated more FNP-59 than vehicle treated AT-3 tumors, reproducing results with fluorescent cholesterol analogs (Figure 5B). Furthermore, T cells recovered from the anti-PD1 treated EO771 tumors had significantly greater uptake of FNP-59 than anti-PD1 treated AT-3 tumors (Figure 5C). These experiments demonstrate that baseline uptake of FNP-59 before ICI treatment corelates with response to therapy in these models; and 2) change in uptake of this radioactive cholesterol analog reflects T cell response to ICI.

**Figure 5.**
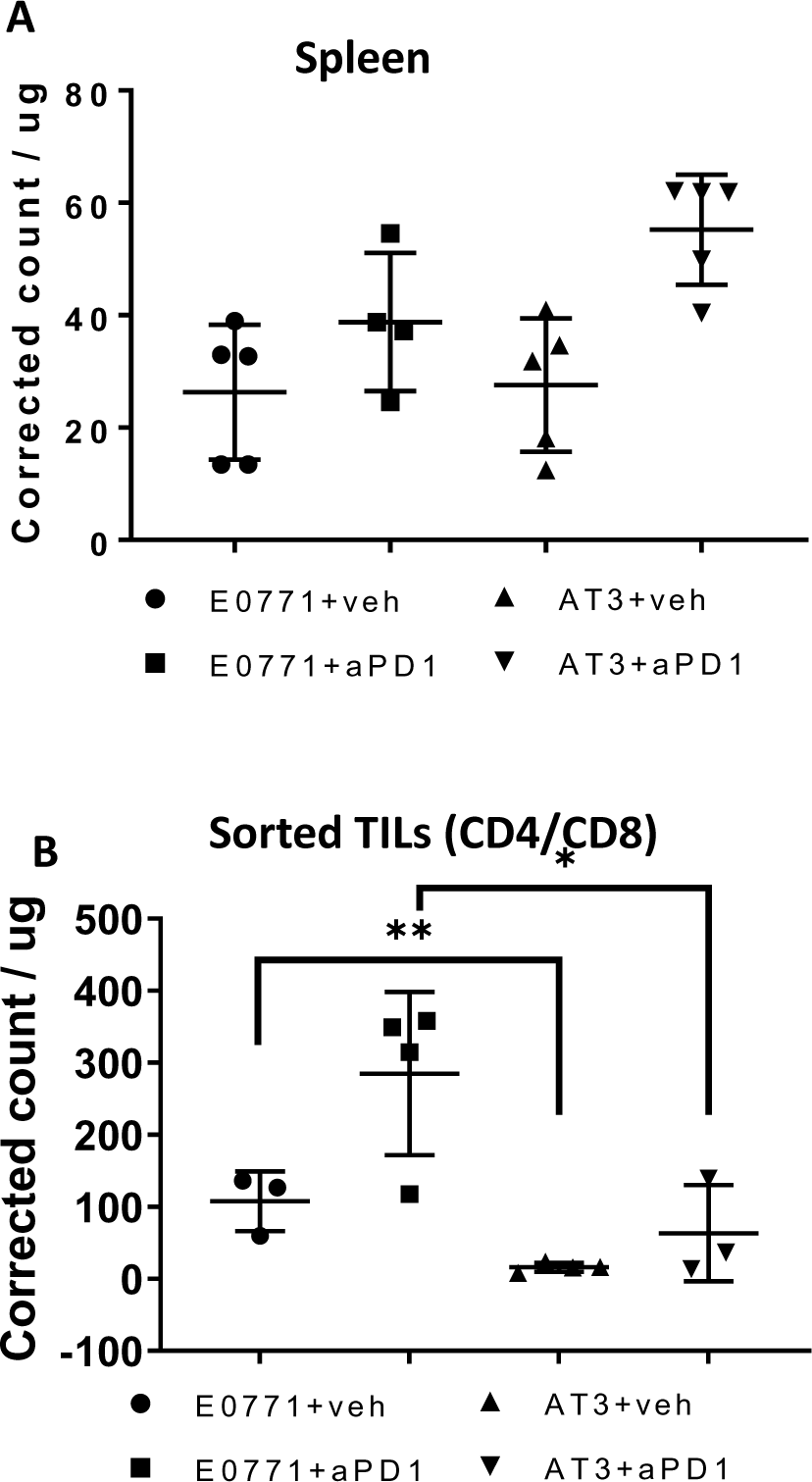
T cells in ICI responsive tumors had greater uptake of FNP-59. When tumors reached ∼ 70 mm^2^, we treated mice with anti-PD-1 antibody or isotype/vehicle for 4 days. We then injected mice with 100 µCi of FNP-59 to quantify uptake in T cells. A) Graph shows uptake of FNP-59 per microgram of spleen tissue measured by scintigraphy. Symbols show individual mice with annotations for mean values and standard deviations. (B) We isolated CD4 and CD8 tumor infiltrating lymphocytes (TIL) by positive selection with immunomagnetic beads and determined accumulation of FNP-59 normalized to total cell protein with BCA assay. Representative data from 2 experimental replicates with statistical comparisons by Student’s T-test.

### Cycling T cells in human TNBC tumors maintain elevated expression of cholesterol trafficking genes

To translate these discoveries to human breast cancer, we queried publicly available datasets and analyzed single cell RNA sequencing data from human TNBC (30). Louvain clusters of TNBC patient cells, without additional characterization or bias, reported four major T cell states (Figure 6A). These states included resting/naïve T cells, highly activated/cycling T cells, transitional effectors (toward exhaustion), and exhausted effectors. We annotated T cell states based on highly expressed genes in each Louvain cluster: 1) Cycling T cells are defined by the high expression of genes related to microtubule reorganization and mitosis including upregulation of STNM1 and MKI67; 2) Transitional state cytotoxic T cells based on high expression of GZMK and moderate levels of PDCD1 and HAVRC2; 3) Dysfunctional T cells classified by high expression of PDCD1, HAVCR2, CXCL13, and expression of CSF1; and 4) Resting memory T cells defined by high expression of CXCR4 and high expression of ribosomal proteins(31–33). Cell counts of clusters from each patient revealed that patients with large T effector populations (whether transitional or exhausted) also carried a smaller population of actively proliferating T cells (Figure 6B). We determined and annotated T cell states of each cluster using top gene expression of the Louvain clusters (figure 4B). Cycling T cells displayed the greatest expression of genes directly and indirectly involved in cholesterol uptake and trafficking (figure 6C). Furthermore, genes involved in cholesterol distribution (Figure 6D), endosomal transport (Figure 6E&F), receptor recycling (Figure 6G, receptor expression (Figure 6I), and transcription factor control (Figure 6I) were significantly modulated in the Cycling T cell populations when compared to other clusters (Supplemental Table 1 for all comparisons). When analyzed using Enrichr we found families of genes related to “Regulation Of Cholesterol Biosynthesis By SREBF” and “Activation Of Gene Expression By SREBF” are significantly expressed in the ‘cycling’ T cell cluster (Supplemental Table 2). Overall, these data show some patients with TNBC, develop tumors that produce a population of cycling T cells (often accompanied by an expanded population of effector T cells; Figure 6B), with features of gene expression indicative of cholesterol uptake and trafficking. Together, these data suggest that increased uptake of cholesterol demarks activated, anti-tumor T cells in human TNBC.

**Figure 6.**
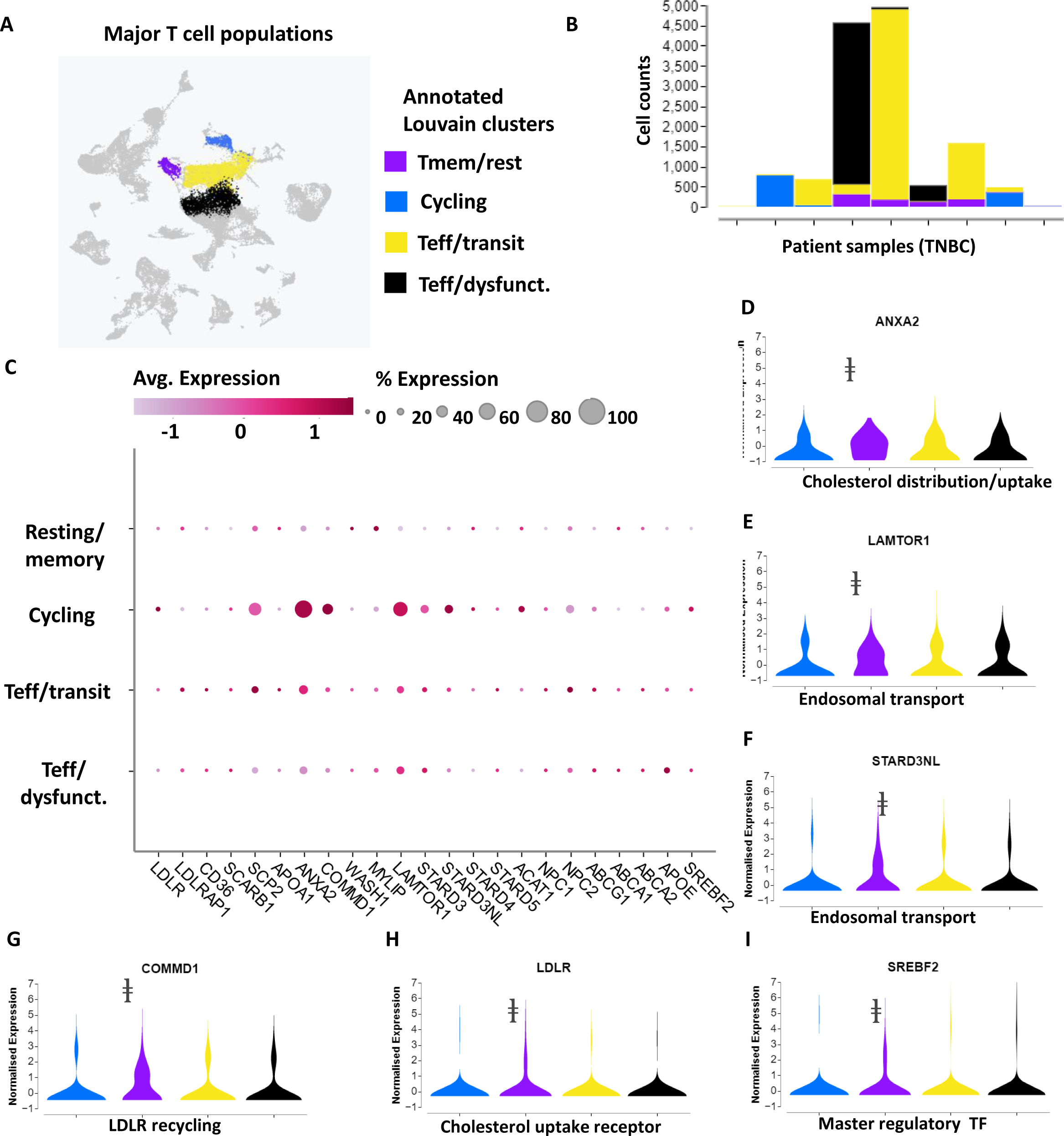
Cycling T cell populations in patients with triple-negative breast cancer upregulate genes related to cholesterol metabolism. (A) Re-analysis of single-cell RNA sequencing data [29] with Cellenics software displays Louvain clusters of annotated T cell states: T memory/resting, highly activated, T effector/transitional state, and T effector/dysfunctional. B) Plot shows proportions of the absolute counts across for various T cell subsets across TNBC patients (n =8429 cells from 9 tumors). C) Clustered averaged gene expression data reveal upregulation of relevant genes involved in uptake and intracellular trafficking of cholesterol in cycling T cells. (D-I) Violin plots of normalized expression of specific genes: D) ANXA2 (cholesterol distribution to the plasma membrane), (E) LAMTOR1 (endosomal transport), (F) STARD3NL (endosomal transport), (G) COMMD1 (LDLR recycling), (H) LDLR (cholesterol uptake), (I) SREBF2 (positive regulation of cholesterol uptake and synthesis). **ⱡ** Indicates P < 10^-20^ between “cycling” cluster and other T cell clusters.

## Discussion

Tumors can generally present as a 1) immune desert phenotype lacking immune cells and immune signaling, 2) an immune excluded phenotype in which immune cells encounter mechanical and cell signaling barriers preventing tumor infiltration, and 3) immune infiltrated or inflamed phenotypes(34). TNBC in humans is heterogeneous, but presents with an inflamed microenvironment more frequently than other subtypes of breast cancer. Although generally considered more inflamed, TNBC can exist anywhere on this continuum of immune phenotypes and exhibits substantial intra- and intertumoral heterogeneity. Tumors with high infiltration of lymphocytes such as NK cells, cytotoxic T cells, and B cells correlates with more favorable prognosis(35). The key publication from Hu et. al, 2021 that has guided immunotherapy clinical trials and practice in TNBC identified three major immune subtypes. Only the subtype with enrichment of lymphocytes and immune checkpoint ligands showed favorable outcomes (36). However, even the most favorable subtype of TNBC demonstrated heterogeneity in TIL function and overall efficacy (through modulation of key genes). Clinical trials and treatment with immunotherapy rely on a combined positive score (CPS) to subclassify patients and predict response. The CPS system analyzes numbers of PD-L1 (or other marker) positive tumor cells, lymphocytes, and macrophages out of total number of viable cancer cells in histologic sections from biopsies. While used widely, the CPS may vary based on selection of different antibodies for immunostaining, and CPS cut-offs have only modest success for predicting response to ICIs(37). Therefore, identification of more informative and potentially non-invasive biomarkers for efficacy of ICIs remains a highly active area of translational research(38).

Given past evidence linking T cell activation with increased cholesterol, we investigated uptake of cholesterol as a novel marker for efficacy of immunotherapy in mouse models of TNBC. We discovered that increasing levels of TCR activation correlated with higher uptake of cholesterol in T cells. Capitalizing on this observation, we leveraged FNP-59, a validated radiotracer for imaging cholesterol trafficking and uptake in humans, as a marker for efficacy of ICI. For tumor studies, we utilized two mouse models of TNBC, EO771 and AT-3, that are responsive or not responsive to immunotherapy, respectively. Ex vivo restimulated and in vivo T cells from ICI responsive breast tumors in mice accumulated significantly more cholesterol than T cells from ICI non-responsive tumors. Remarkably, we detected uptake of FNP-59 within three hours of injection in tumor-bearing mice. T cells from tumors responsive to ICI therapy showed higher uptake of FNP-59 under basal conditions and following anti-PD1 therapy. In human patients with TNBC, populations of “activated T cells” exhibited gene signatures of enhanced cholesterol uptake as compared with resting, effector, and exhausted T cell states. This gene signature included upregulation of key genes, including ANXA2, a multifaceted protein that works with CD36 for lipid scavenging(39). Additionally, we found upregulation of the master transcriptional regulator of cholesterol metabolism, SREBF2(40).

Predicting which patients will benefit from immunotherapy remains a central obstacle to effective use of these drugs for cancer. There are currently three FDA-approved predictive markers for immunotherapy: PD-L1 expression, tumor mutational burden (TMB), and microsatellite instability (MSI). Approved methods rely on tumor biopsies, limiting the ability to repeatedly monitor tumor evolution and response to therapy over time. Several other investigational approaches exist, including gene profiles for T cells and presence of B cells(41). Listed strategies focus on predicting response to immunotherapy to guide selection of patients. However, none of these methods offer the important capability to detect response to therapy early in treatment.

Toward the goals of predicting response to ICIs and detecting early effects of therapy, researchers have developed PET imaging probes based on antibodies, antibody fragments such as affibodies or small molecules for the PD-L1/PD-1 pathway or other immune checkpoint inhibitors(42, 43). PET imaging studies in mouse models or early clinical studies reveal heterogeneous expression of immune checkpoint proteins in primary tumors and various metastases, emphasizing the value of imaging to detect intra-patient heterogeneity that could affect treatment outcomes(44). As discussed previously, detecting expression of PD-L1 protein or other immune checkpoint molecules at least by immunostaining has shown limited efficacy to predict patients who will respond to ICIs. Imaging probes to cell surface immune checkpoint inhibitors identifies levels of specific targets, and imaging after starting therapy can detect to what extent an ICI blocks the target molecule. A key limitation is that these probes do not detect to what extent a therapy activates functional T cells, the key effector population of current ICIs.

Functions of cholesterol in T cells and anti-tumor immunity remain controversial. One study found that hyperlipidemia and increased cholesterol plasma disrupted T cell homeostasis and drove expansion of effector memory T cells(19). In contrast, Xingzhe Ma, et al found that cholesterol in the tumor microenvironment correlated with T cell exhaustion due to ER stress(6). An alternative or additional hypothesis is that the presence of exhausted T cells indicates a prior active immune response in those tumors. Notably, the study by Ma et al did not show to what extent the effect of cholesterol on T cells reversed with immunotherapy. In our study we found greater success of ICI therapy in breast tumors wherein T cells accumulated greater cholesterol from the extracellular environment. A recent study comprehensively mapped cholesterol metabolism and disposition in tumor environments(45). Intratumoral T cells showed cholesterol deficiency, which impaired proliferation and triggered autophagy-mediated cell death. Across all cells in a tumor, tumor cells and myeloid lineage cells contained highest amounts of cholesterol with lower overall levels in T cells. Intriguingly, suppressing cholesterol efflux pathways by targeting the LXRβ transcription factor enhanced anti-tumor functions of T cells. These data support our findings that activated T cells accumulate more cholesterol and point to the need for methods to track disposition of cholesterol in tumor environments. Methods used in this study focused on uptake of cholesterol in relation to effects of immunotherapy. Our data do not address to what extent, if any, overall increases in cholesterol or cholesterol metabolism in T cells regulate antitumor immunity.

FNP-59 has limitations for PET imaging of ICIs in cancer because uptake of the radiotracer and cholesterol in a tumor is not restricted to T cells. Signal from FNP-59 would also include uptake of cholesterol by macrophage, cancer cells, and other cell types. Potentially, measuring an increase in FNP-59 uptake in a tumor between imaging studies conducted before and shortly after starting ICI treatment could reflect accumulation in re-activated T cells. This observation could be confounded by recruitment of more T cells or other immune cell types (also increasing uptake of FNP-59) in the short term, although such changes would be consistent with a more inflamed tumor environment. From our study we can conclude that ex vivo analysis of this PET radiotracer could be used to measure redistribution of cholesterol in a tumor in pre-clinical models of biopsy samples.

FNP-59 presents numerous benefits over the previous iodinated version of the compound. The poor safety profile, onerous synthetic protocols, and poor image quality restricted the iodinated version (131I-NP59) of this tracer to detecting diseases of the adrenal cortex. Use of 131I-NP59 has been discontinued in the United States [42]. FNP-59 overcomes many of these limitations by the previous generations of this tracer, and initial studies in humans show ideal imaging characteristics and no toxicity. These studies provide a roadmap for potential clinical translation of this radiotracer for monitoring treatment with ICIs in a wide variety of cancers.

Overall, efficacy of FNP-59 for identifying activated, anti-tumor immune cells in mice, combined with relevance of cholesterol uptake in activated T cells in humans, support future testing of this radiotracer to stratify patients and monitor response to ICI in TNBC and possibly other malignancies. This radiotracer could be used in TNBC before biopsy or surgery, followed by isolation of T cells and ex vivo imaging of accumulation. Alternatively, since early responses to ICI therapy include activation of peripheral blood CD8 T cells enriched for anti-tumor T cell clones, changes in accumulation of radiotracer in blood samples collected before and after treatment could monitor efficacy of these drugs. Future studies will investigate these potential applications of FNP-59 for prediction and treatment monitoring with ICIs in mouse models of breast cancer and ultimately patients.

## Methods

### Cell lines

We cultured mouse EO771 (ATCC) and AT-3 (EMD Millipore) breast cancer cells and C57BL/6J mouse mammary fibroblasts (gift of Harold Moses, Vanderbilt University) in DMEM base media (Gibco, #11965) supplemented with 10% FBS, 1% GlutaMax (ThermoFisher Scientific, #35050061), and 1% penicillin-streptomycin (Gibco, #15140148). We maintained all cells in a humidified 37°C incubator with 5% CO_2_.

### Mouse tumor models

The University of Michigan Institutional Animal Care and Use Committee approved all animal procedures under protocol PRO00010534. We implanted 5×10^5^ EO771 (ATCC) or AT-3 (EMD Millipore) mouse mammary tumor cell lines and 1×10^5^ mouse mammary fibroblasts orthotopically into 4^th^ inguinal mammary fat pads of 6-8-week-old female C57BL/6J female mice (The Jackson Laboratory)(46). We housed mice in specific pathogen-free housing facilities maintained by the Unit for Laboratory Animal Medicine at the University of Michigan. Three days after implanting cells, we randomly assigned mice to treatment with anti-PD-1 antibody (10 mg/kg intraperitoneal) (BioXCell, #CD279) or isotype/vehicle ctrl administered every three days for four total doses. We measured growth of orthotopic tumors with calipers twice per week, quantifying tumor size for each mouse as mm^2^ by multiplying orthogonal length x width measurements. We euthanized mice for a pre-determined humane endpoint of tumor size, which defined time of censoring for survival analyses.

### T cell assay preparation, reagents, and antibodies

We purified T cells from the spleens of wild-type C57BL/6J mice by processing tissue through a 70-micron filter, lysing red blood cells with ACK buffer (vendor and number), and sorting cells using the Pan T Cell Isolation Kit II from Miltenyi (#130-095-130). For ex vivo activation of T cells, we coated 96 well plates with indicated amounts of anti-CD3ε antibody supplemented with soluble anti-CD28 at 2-times the amount of anti-CD3ε and 5U/mL hL IL-2 (Biolegend). For activation, specific antibodies included anti-mouse CD3ε (145-2C11) and CD28 (37.51) or anti-human CD3 antibody (OKT3) and anti-human CD28 (CD28.2) (antibodies from Biolegend). For flow cytometry experiments, antibodies included anti-mouse CD45(S18009D), TCRβ chain (H57-597), CD3 (17A2), CD8α (53-6.7), CD4 (RM4-5), CD69 (H1.2F3), and CD279 (29F.1A12) (Biolegend). Cholesterol probes included Cholesteryl BODIPY 542/563 C11 (Invitrogen, #C12680) and 3-dodecanoyl-NBD Cholesterol (Cayman Chemicals, #13220) for in vivo and in vitro experiments respectively.

### Synthesis of FNP-59

#### The synthesis of FNP-59 was accomplished as previously described for clinical research studies

[^18^F]Fluoride was produced via the ^18^O(p,n)^18^F nuclear reaction with a GE PETtrace cyclotron equipped with a high-yield fluorine-18 target. [^18^F]Fluoride was delivered in a bolus of [^18^O]H_2_O to the synthesis module and trapped on a QMA-Light sep-pak cartridge to remove [^18^O]H_2_O. [^18^F]Fluoride was then eluted into the reaction vessel with potassium carbonate. Acetonitrile with Kyrptofix 2.2.2 was added to the reaction vessel, and the [^18^F]fluoride was azeotropically dried by heating the reaction vessel to 100 °C and drawing full vacuum. After this time, the reaction vessel was subjected to both an argon stream and a simultaneous vacuum draw at 100 °C. The solution of precursor in DMSO was added to the dried [^18^F]fluoride, and it was heated at 120 °C with stirring for 20 min. Ester group was removed by treatment with KOH and heating at 110 °C for 25. FNP-59 was isolated via reverse phase chromatography and reformulated into a solution of saline, ethanol and Tween-80.

### In vitro/In vivo uptake of FNP-59

For experiments measuring T cell accumulation of FNP-59 in vitro, we washed 96 well plates with activated cells once before adding radiotracer at indicated concentrations (n = 3 wells per concentration). We incubated plates for 1 hour and then washed twice with RPMI medium (Gibco, #11875903) containing 0.1% serum before quantifying retained radioactivity in each well. We performed all radioactivity measurements for experimental wells and a standard curve with a phosphor imaging screen (GE Typhoon FLA 7000). Image analysis was performed using ImageQuant (Molecular Dynamics) software. Regions-of-interest were drawn and converted to disintegrations per minute (DPM) and specific activity using molecular standards on the same plate and data was further modeled and plotted with Prism GraphPad version 9 (San Diego, CA, USA).

For mouse experiments, we intravenously injected 100 µCi of FNP-59 three hours before imaging selected mice or euthanizing mice to recover spleen and tumor-infiltrating T cells. We euthanized all other mice three hours after injection. To recover T cells for analysis, we dissociated tumors in 5mL of tumor dissociation buffer containing RPMI base media, 5% FBS, 300U/mL Collagenase I, 150U/mL Collagenase IV and, 200ug/mL of DNase1 (VENDORS), using the TDK1 program on the gentleMACS Octo Dissociator (Miltenyi, Waltham, MA, USA). After 30 minutes of incubation, we passed tumor tissue through a 70-micron filter and isolated T cells using CD4/CD8 (TIL) MicroBeads and cell sorting magnets from Miltenyi. Recovered T cells were placed into 1.5mL tubes and measured decay-corrected radioactive counts with a (gamma) well counter. We quantified accumulation of FNP-59 in excised whole spleens. We normalized accumulated radiotracer activity to total protein via BCA assay (Pierce assay) for T cells and whole organ weight for the spleen, respectively.

### In vitro/ In vivo fluorescent cholesterol uptake and flow/spectral cytometry

For in vitro experiments measuring uptake of fluorescent cholesterol analogs during early T cell activation, we added 1ug/mL 3-NBD cholesterol to 96 well plates with T cell activation conditions for 24 hours. We washed T cells once with PBS and prior to staining in the plate using antibodies against T cell identification markers (TCRβ, CD4, and CD8) and activation marker CD69. We analyzed cells on the same day using a Bio-Rad Ze5 Analyzer (Bio-Rad, Hercules, CA, USA). Single color controls enabled compensation and spectral unmixing of samples, and fluorescence minus one (FMO) control ensured no autofluorescence or incidental FRET with emission from BODIPY. We used the following gating strategy: 1) FSC/SSC to exclude debris; 2) single cells defined by FSC-H/FSC-A gated on the linear population; 3) live cell populations using live-dead exclusion dye (Zombie Aqua, #423101, Bio-Rad); 4) CD45+ to include only white blood cells; 5) TCRβ constant chain-positive to include all T cells; and 6) CD4 or CD8 gates and finally analytic antibodies or probes including CD69, PD-1, or cholesterol.

To analyze total T cell cholesterol content, we used the Cholesterol/Cholesterol Ester-Glo™ Assay from Promega. The experiments were setup in a 96 well plate with 10^5^ cells per well. Cells were stimulated with plate bound anti-CD3 using 0.01, 0.1, 1, and 10 μg/mL and an additional group was unstimulated. Soluble anti-CD28 was provided to all groups at three times the amount of anti-CD3 for each group and IL-2 was excluded to prevent the impact of rapid growth stimulation and dissociate the effect from added growth factor/high affinity IL-2 receptor expression. After 24 hours, samples were diluted in Cholesterol Lysis Solution and the manufacturer’s procedure was followed with 1:8 and 1:20 dilutions of sample. This protocol measures total cholesterol with samples assayed plus esterase in Cholesterol Detection Reagent and free cholesterol with samples assayed minus esterase in Cholesterol Detection Reagent. Cholesterol ester levels are calculated as the difference between total and free cholesterol. Luminescence was recorded on an IVIS Lumina LT series III (Perkin Elmer).

For in vivo experiments, we injected mice intraperitoneally with 150 µg of BODIPY labeled cholesterol in 50% DMSO + PBS (added directly before injection) or vehicle only. After 1 day, we euthanized mice; removed tumors; and weighed each tumor. We dissociated tumors as described for radiotracer studies and used 10^6^ cells for staining for flow cytometry.

### Data mining of scRNA-seq data

To reanalyze data from Wu SZ et al. 2021, we downloaded sequencing files from the Single Cell Portal hosted by the Broad institute at https://singlecell.broadinstitute.org/single_cell/study/SCP1039/a-single-cell-and-spatially-resolved-atlas-of-human-breast-cancers#study-download (brca_mini_atlas_raw_unfiltered.zip). We analyzed the dataset with Cellenics software (Biomage). Analysis included reprocessing the data by classifier filter, mitochondrial content filter, GenesVsUMI filter, doublet filter, data integration with Seurat v4, and UMAP embedding. Data processing settings can be found in the supplemental file “*data processing settings.txt*” with data found in supplemental file “*processed_matrix.rds*”. Cellenics web user interface provided further options for data visualization including UMAP, cluster cell counts, DOT plots and individual gene violin plots.

To analyze data from, Martin, et. al. 2019, we used the web user interface from The Broad Institute at, https://singlecell.broadinstitute.org/single_cell/study/SCP359/ica-ileum-lamina-propria-immunocytes-sinai?scpbr=immune-cell-atlas#study-visualize as directed from the Immgen database at https://www.immgen.org/Databrowser19/HumanExpressionData.html. We retrieved data presented in this paper from the web user interface and compared the selected genes between activated T cells and other effector populations. Data can be retrieved and replicated through the above link.

### Statistics

We performed all data processing in Prism GraphPad V9. Student’s t test was used to compare statistical significance between groups of normally distributed samples and mean fluorescence intensity, or relative means were analyzed using nonparametric Mann-Whitney tests. Asterisks (ns = p > 0.05; *p < 0.05, **p < 0.01,***p < 0.001, ****p < 0.0001) indicate the degree and magnitude of significance between two groups. We created Kaplan–Meier plots and analyzed differences in survival using the Gehan–Breslow–Wilcoxon test. In assessing statistical differences of tumor burden over time between in vivo groups, we used non-linear regression for logistical growth to determine significant differences between growth curves. Similarly, for in vitro studies using FNP-59, non-linear regression and tests between data set parameters were used to assess significance. A p-value cutoff of p < 10^-20^ defined significant differences in expression of specific genes between Louvain clusters.

## Author contributions

GDL, AFB, NGC, and BV conceptualized this study. NGC and JG analyzed the data and NGC wrote the manuscript. Primary drafting of the manuscript by NGC (listed first) with essential editing and input by GDL, BV, and AFB. NGC, JG, & AF performed in vitro/ex vivo T cell studies. NGC, AF, & JG performed all in vivo experiments. NGC solely performed data query, mining, and re-analysis. NGC,TD&JS performed all in vivo and in vitro Radiation/PET tracer experiments using the laboratory and resources of AFB. NGC, AFB, BV, PS, and GDL contributed to the written discussion and scientific discussions around the manuscript that lead to its drafting and formation.

## Acknowledgments

The authors acknowledge funding from United States National Institute of Health grants R01CA238042, R01CA238023, R33CA225549, U24CA237683, and R37CA222563 (G.D.L). We also acknowledge support from the United States Department of Defense Breast Cancer Research program award W81XWH2210120 and from the W.M. Keck Foundation (G.D.L). Funding for radiotracers came for Fast Forward Medical Innovation MTrac, Michigan Memorial Phoenix Project, and the Department of Radiology (B.L.V.). The manuscript includes data generated through Flow Cytometry, and Pre-Clinical Imaging shared resources supported in part by the University of Michigan Comprehensive Cancer Center support grant P30CA046592.

