## Supplemental figures for "Predicting efficacy of immunotherapy in mice with triple negative breast cancer using a cholesterol PET radiotracer"

### Supplemental figure 1

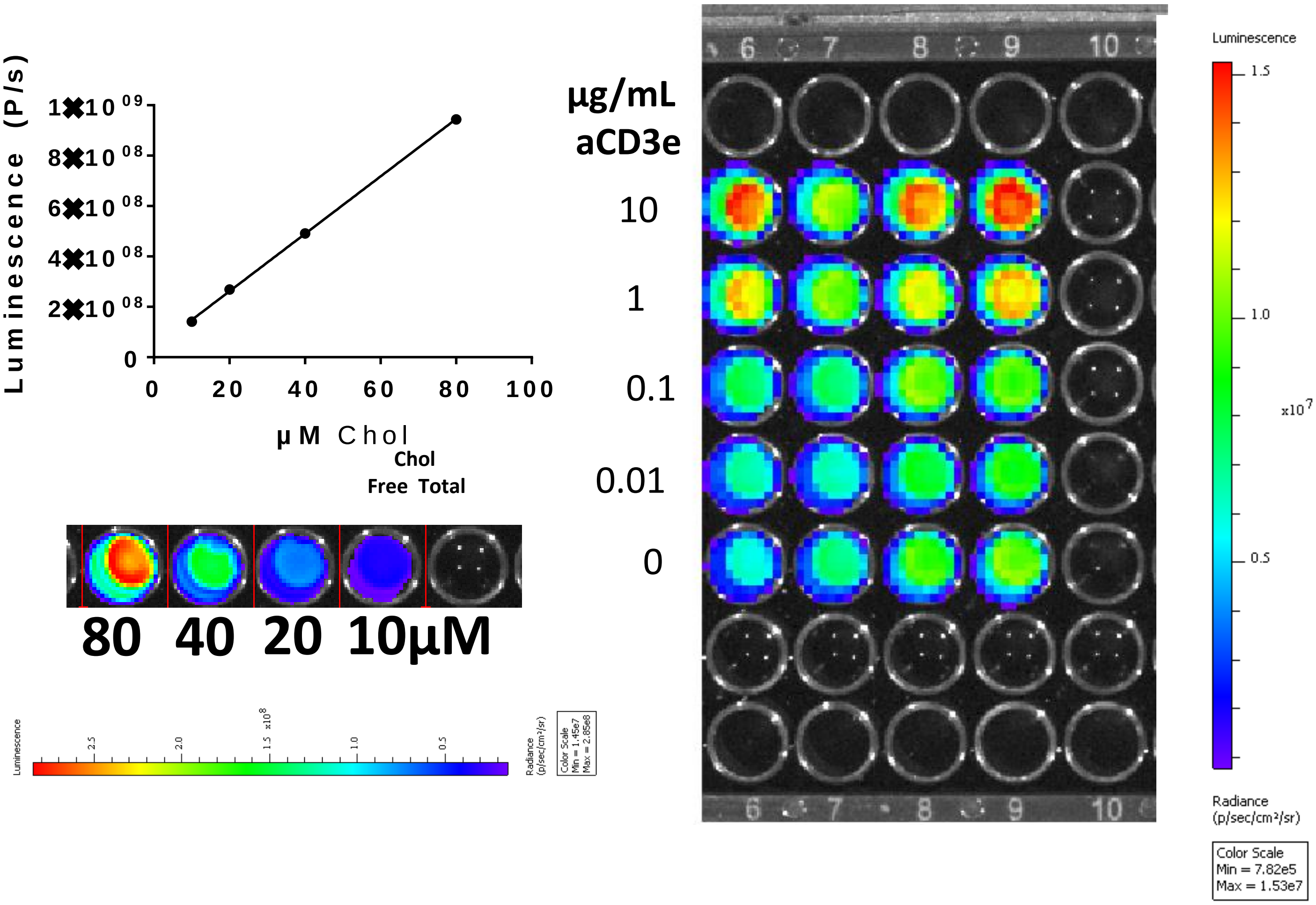


**Supplemental Figure 1.** Standard curve and luminescence image for luminescent cholesterol assay shown in figure 4A. Plate image after IVIS imaging. Anti-CD3 concentrations were plated decreasing in rows top to bottom as labeled and in duplicate with esterase added on the right (for total cholesterol).

### Supplemental figure 2

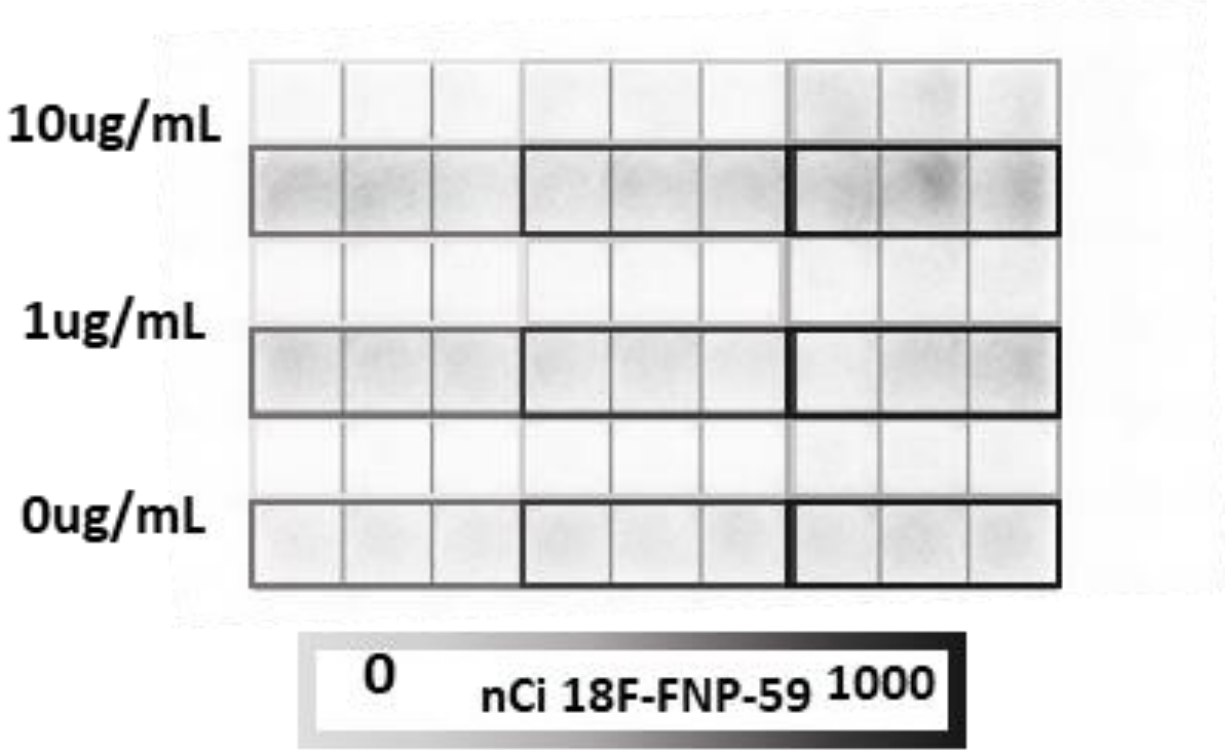


**Supplemental Figure 2.** Example of plate image after phosphor screen processing (Figure 4C). FNP-59 concentrations were plated increasing left to right in triplicate. The six concentrations represented by light gray to black color boxes were 0.05, 0.1, 0.25, 0.5, 0.75, 1 µCi treatments over 1 hour. For example, row one has 0.05, 0.1 and 0.25 µCi treatment (left to right) while row 2 has 0.5, 0.75, and 1 µCi treatments and these top 2 rows were stimulated with 10ug/mL anti-CD3.

### Supplemental Table 1

| pAdj values (fig 6D-I) | | | | Log FC expression (fig 6D-I) | | | |
|---|---|---|---|---|---|---|---|
| **Cycling vs** | **Rest /mem** | **Transit /eff** | **Dysfunct.** | | **Rest /mem** | **Transit /eff** | **Dysfunct.** |
| ANXA2 | 1.68E-68 | 4.68E-48 | 2.29E-70 | ANXA2 | 0.6526 | 0.4209 | 0.5317 |
| SREBF2 | 3.56E-26 | 6.74E-34 | 3.36E-26 | SREBF2 | 0.09925 | 0.06603 | 0.05973 |
| LAMTOR1 | 1.11E-51 | 3.87E-36 | 1.10E-20 | LAMTOR 1 | 0.40000 | 0.2548 | 0.2007 |
| LDLR | 1.23E-20 | 9.92E-37 | 2.08E-56 | LDLR | 0.1059 | 0.09252 | 0.1155 |
| COMMD1 | 2.11E-52 | 1.27E-59 | 4.43E-55 | COMMD 1 | 0.04908 | 0.1958 | 0.2055 |
| STARD3NL | 4.87E-45 | 6.40E-46 | 7.69E-53 | STARD3N L | 0.2214 | 0.1502 | 0.1787 |
| **Rest/mem vs.** | | **Transit /eff** | **Dysfunct.** | | | **Transit /eff** | **Dysfunct** |
| ANXA2 | | 1.385e-13 | 1.216e-5 | ANXA2 | | -0.2317 | -0.1209 |
| SREBF2 | | 0.001272 | 0.1003 | SREBF2 | | -0.03322 | -0.03952 |
| LAMTOR1 | | 8.771e-9 | 3.014e-14 | LAMTOR 1 | | -0.1441 | -0.1983 |
| LDLR | | 0.1635 | 0.9657 | LDLR | | -0.01337 | 0.009653 |
| COMMD1 | | 6.231e-5 | 2.402e-4 | COMMD 1 | | -0.06401 | -0.05425 |
| STARD3NL | | 2.552e-6 | 1.165e-3 | STARD3N L | | -0.07114 | -0.04269 |
| **Teff vs.** | | | **Dysfunct** | | | | **Dysfunct** |
| ANXA2 | | | 6.498e-7 | ANXA2 | | | 0.1107 |
| SREBF2 | | | 1 | SREBF2 | | | -0.006304 |
| LAMTOR1 | | | 0.002409 | LAMTOR 1 | | | -0.05415 |
| LDLR | | | 2.394e-5 | LDLR | | | 0.02302 |
| COMMD1 | | | 0.4722 | COMMD 1 | | | 0.009758 |
| STARD3NL | | | 0.07073 | STARD3N L | | | 0.02845 |

Supplemental Table 1. We re-analyzed major T cell populations from existing single-cell RNA sequencing data [29] . Adjusted P-values and logFC values of specific genes expressed when comparing Louvain clusters from Figure 6. Rest/mem is the annotated resting/memory T cell population, Transit/eff is the transitional/effector T cell population, and Dysfunct is the dysfunctional T cell population.

### Supplemental Table 2

| **Cycling T cells Vs** | | | | |
|---|---|---|---|---|
| Name | P-value | Adj P-value | Odds ratio | Combined score |
| **Regulation Of Cholesterol Biosynthesis By SREBP (SREBF) R-HSA-1655829** | **7.886e-8** | **9.261e-7** | **8.28** | **135.43** |
| **Activation Of Gene Expression By SREBF (SREBP) R-HSA-2426168** | **0.000002717** | **0.00002195** | **8.44** | **108.11** |

**Supplemental Table 2. Increase expression of genes related to cholesterol uptake in cycling T cell populations in patients with triple-negative breast cancer**. We re-analyzed major T cell populations from existing single-cell RNA sequencing data [29] using Cellenics software by Biomage (see Figure 6). With Reactome analysis software interface, we compared the "cycling" cluster to all other T cells using Enrichr for pathways ontologically related to cholesterol uptake.  Filter cutoffs used P < 0.05 and logFC > 0.1.
